# Met-ID: An Open-Source Software for Comprehensive Annotation of Multiple On-Tissue Chemical Modifications in MALDI-MSI

**DOI:** 10.1101/2025.01.24.634674

**Authors:** Patrik Bjärterot, Anna Nilsson, Reza Shariatgorji, Theodosia Vallianatou, Ibrahim Kaya, Per Svenningsson, Lukas Käll, Per E. Andrén

## Abstract

Here, we introduce Met-ID, a graphical interface software designed to efficiently identify metabolites from MALDI-MSI datasets. Met-ID enables annotation of *m/z* features from any type of MALDI-MSI experiment, involving either derivatizing or conventional matrices. It utilizes structural information for derivatizing matrices to generate a subset of targets that contain only functional groups specific to the derivatization agent. The software is able to identify multiple derivatization sites on the same molecule, facilitating identification of the derivatized compound. This ability is exemplified by FMP-10, a reactive matrix that assists the covalent charge-tagging of molecules containing phenolic hydroxyl and/or primary or secondary amine groups. Met-ID also permits users to recalibrate data with known *m/z* ratios, boosting confidence in mass match results. Furthermore, Met-ID includes a database featuring MS2 spectra of numerous chemical standards, consisting of neurotransmitters and metabolites derivatized with FMP-10, alongside peaks for FMP-10 itself, all accessible directly through the software. The MS2 spectral database supports user-uploaded spectra and enables comparison of these spectra with user-provided tissue MS2 spectra for similarity assessment. Although initially installed with basic data, Met-ID is designed to be customizable, encouraging users to tailor the software to their specific needs. While several MSI-oriented software solutions exist, Met-ID stands out as a pioneer in seamlessly combining both MS1 and MS2 functionalities. Developed in alignment with the FAIR Guiding Principles for scientific software, Met-ID is freely available as an open-source tool on GitHub, ensuring wide accessibility and collaboration.

## INTRODUCTION

Metabolomic analysis using mass spectrometry imaging (MSI), through matrix-assisted laser desorption/ionization (MALDI), has emerged as a crucial tool for studying the spatial distribution of metabolites in biological tissues^1-3^, providing insights into the metabolic heterogeneity of tissues. When applied to biological tissues, MALDI-MSI can produce hundreds or even thousands of mass-to-charge (*m/z*) features in a single experiment^4^. However, identification of metabolites in these studies may be challenging due to the vast chemical diversity and low ionization efficiency of many metabolites. Some metabolites ionize poorly during MALDI-MSI, resulting in weak or undetectable signals, which limit their detection. Results may be further complicated by ion suppression from more easily ionized compounds, making it difficult to identify less ionizable metabolites accurately^5^.

A prominent addition to the technique that enhances the detection of diverse metabolites is on-tissue chemical derivatization^5, 6^. This modifies metabolites directly on tissue sections, introducing permanent charges to the structure of target molecules and enhancing their ionization characteristics and detectability in MALDI-MSI. Derivatizing agents can be used as complements to conventional matrices or on their own as reactive matrices, depending on which metabolites are to be studied^7^. There are many examples of reactive matrices and derivatizing agents in the literature, e.g., FMP-10 (4- (anthracen-9-yl)2-fluoro-1-methylpyridin-1-ium iodide)^5^, TAHS (*p*-*N,N,N*-trimethylammonioanilyl *N*-hydroxysuccinimidyl carbamate iodide)^8^ and Girard-T^9^.

The reactive matrix FMP-10 was originally developed for the analysis of neurotransmitters, including those within the dopaminergic and serotonergic systems, and their related metabolites^5^. It specifically targets metabolites that contain primary amines and phenolic hydroxyl groups, covering a comprehensive range of compounds within the dopaminergic and serotonergic pathways, in addition to various amino acids and other metabolites. Unlike traditional matrices that assist in the ionization process, FMP-10 and similar derivatizing matrices possess permanent charges. Upon forming covalent bonds with target molecules, these charges render the resultant derivatives highly suitable for mass spectrometry analysis. This approach deviates from the conventional measurement of protonated or de-protonated species because the derivatizing matrices generate unique derivatives, each with distinct delta masses. Furthermore, the FMP-10 matrix can derivatize target molecules that have multiple functional groups, leading to formation of several potential ions. For instance, dopamine can undergo single, double or triple derivatization, generating an intricate spectral profile^5^.

Several tools have been developed to annotate metabolites for mass spectrometry. Metlin^10^ and GNPS (Global Natural Products Social Molecular Networking)^11^ are widely used databases for LC-MS research. For MSI, the most commonly used software is Metaspace, which represents a significant advancement in metabolite annotation^12^. This cloud-based platform has become a valuable tool for metabolomic researchers, including those focused on spatial metabolomics, enabling them to upload and analyze datasets. However, Metaspace only provides limited support for annotation of datasets processed with derivatizing matrices. Hence, there is a need for software capable of handling the more complex MSI annotations required by derivatizing matrices. Such software could significantly enhance the quality of metabolite annotations and identification.

In response to these analytical challenges, we have developed new open-source software, called Met-ID, designed to streamline and enhance the identification process of derivatized metabolites in MALDI-MSI. Met-ID facilitates mass matching of the intact ionized molecule (MS1) through enhanced database queries and provides tandem MS (MS2) data of standards, allowing subsequent detailed analyses. It integrates advanced algorithms to automatically annotate the chemically derivatized metabolites, significantly reducing the time and complexity involved in data analysis. The software supports high-throughput metabolomic studies and minimizes potential errors and biases associated with manual data interpretation. By providing a systematic workflow for the annotation and removal of false positives, Met-ID enables researchers to more accurately and efficiently explore the metabolome of biological tissues. Although initially developed for MALDI-MSI, it can be applied to various mass spectrometry-based metabolomic techniques, both spatial and non-spatial. Met-ID is released under an open Apache 2.0 license.

## RESULTS

### Overview of Met-ID

Met-ID has been designed to efficiently handle MS1 and MS2 data in various file formats. For MS1 data, the software supports importing feature lists in .txt and .csv formats, including CSV feature tables generated directly from SCiLS software. The data are displayed in a tabular format, and Met-ID offers two options for exporting results: users can export annotated *m/z* features from the table within Met-ID or generate a comprehensive table of all *m/z* features with adjusted *m/z* values, suitable for downstream analyses. For MS2 data, which is displayed as spectra, Met-ID accepts files in mzML format^13^. Mirrored spectra can be exported either as image files or as CSV files containing the spectral data for further analysis.

While originally developed for Windows, Met-ID has been successfully demonstrated in executable files compatible with MacOS and Linux environments. While Met-ID can process MS1 and MS2 data independently, a typical MSI workflow involves sequential analysis: MS1 followed by MS2 for selected ions (Figure 1).

**Figure 1.**
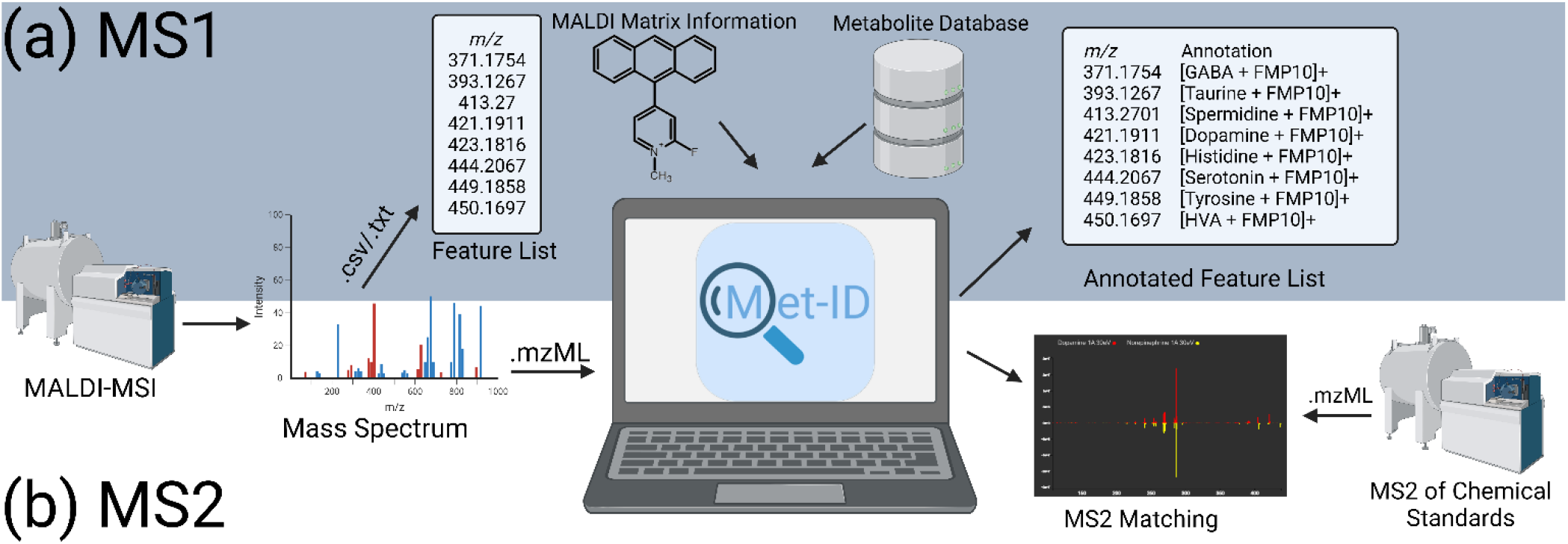
The Met-ID workflow. **(a)** Met-ID processes MS1 data by importing features from a .csv or .txt file, along with MALDI matrix information and a pre-compiled metabolite database, resulting in a list of annotated features for downstream analysis. (b) In the MS2 workflow, Met-ID requires .mzML files of MS2 spectra for comparison with database spectra, highlighting its capability to handle both MS1 and MS2 data.

### Met-ID Database

Met-ID is installed with a database derived from the Human Metabolome Database (HMDB)^14^ and Lipidmaps^15, 16^ and is searchable for common adducts, such as [M+H]^+^, [M+Na]^+^ and [M+K]^+^ in the positive ionization mode and [M-H]^-^, [M+Cl]^-^, [M+Na-2H]^-^, [M+K-2H]^-^ and [M-H_2_O-H]^-^ in the negative ionization mode, along with derivatizing matrices FMP-10^5^ and AMPP (1-(4- (aminomethyl)phenyl)pyridin-1-ium chloride)^17^. Using annotations found in the HMDB and Lipidmaps, the Met-ID database has been constructed in a way to offer users multiple choices. HMDB contains information about the biological origin and organ distribution of metabolites, which allows for higher specificity in Met-ID.

The database also includes MS2 spectra of chemical standards derivatized with FMP-10, allowing graphical comparison with experimental MS2 spectra. To enable Met-ID to be used for any type of MSI experiment, it was developed to be dynamic, i.e., users can add their own metabolites, functional groups and matrices, as well as MS2 spectra to customize their own version of Met-ID. The malleability of Met-ID will be presented at the end of the results section.

### Annotating Chemically Derivatized Metabolites Using Met-ID and MS1

A well-structured approach toward metabolite identification is essential for effective mass spectrometry workflows. Met-ID enhances this process by strategically exploiting the chemical properties of analytes and dynamically adapting database searches (Figure 2). By incorporating a priori knowledge about the specificity of derivatizing matrices, such as FMP-10 and AMPP, Met-ID uses the chemical structures in the form of simplified molecular-input line-entry system (SMILES)^14^ to parse the HMDB and Lipidmaps and more effectively differentiate between potential candidates and non-candidates. Met-ID also allows users to refine their search based on tissue or biofluid type, and whether they are investigating endogenous, exogenous, unspecified or a combination of metabolites.

**Figure 2.**
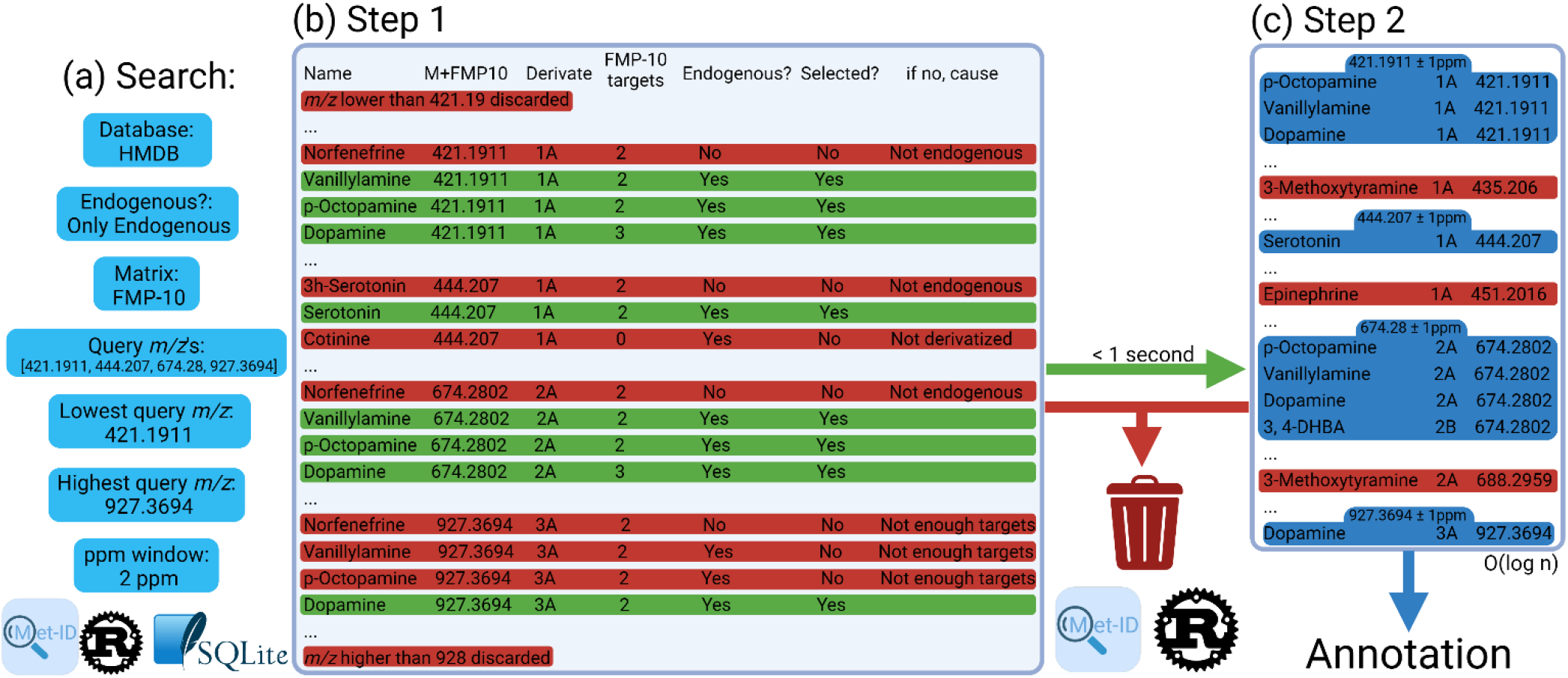
The Met-ID algorithm. (a) The Met-ID search workflow is depicted, showing user-defined search preferences. (b)A table is generated using an SQLite query based on the workflow in (a), aggregating various precompiled Met-ID database tables into a single comprehensive table. From this table, a custom query selects relevant rows. Rows highlighted in red represent metabolites discarded for one or more of the following reasons: the *m/z* ratio falls outside the query range, the specified matrix cannot derivatize the metabolite, insufficient targets are available for a specific derivative, or the metabolite is not endogenous (if this filter is applied). The number of matrix targets is precomputed based on the SMILES structure of each molecule. (c) In Step 2, the green rows retained in Step 1 are further filtered by iterating through the list of query *m/z* values. Using Python’s bisect left algorithm (with a time complexity of O(log n)), the algorithm determines which rows fall within the 2 ppm window of each query *m/z* value. Rows outside the ppm window are marked in red and discarded, leaving the remaining rows in blue as the final annotations displayed in the software.

Met-ID uses chemical information to optimize database queries, aiming to reduce the occurrence of false positives. Figure 3 illustrates four isomers sharing the same formula and monoisotopic mass but exhibiting different molecular structures. FMP-10, which derivatizes primary amines and phenolic hydroxyls (highlighted in blue), does not alter enbucrilate, whereas *p*-octopamine and norfenefrine can potentially undergo two derivatizations and dopamine can undergo three. Consequently, Met-ID selectively queries potential derivatizations excluding compounds like enbucrilate that cannot undergo derivatization, thus avoiding its inclusion in the analysis of the peak under consideration (Figure 3).

**Figure 3.**
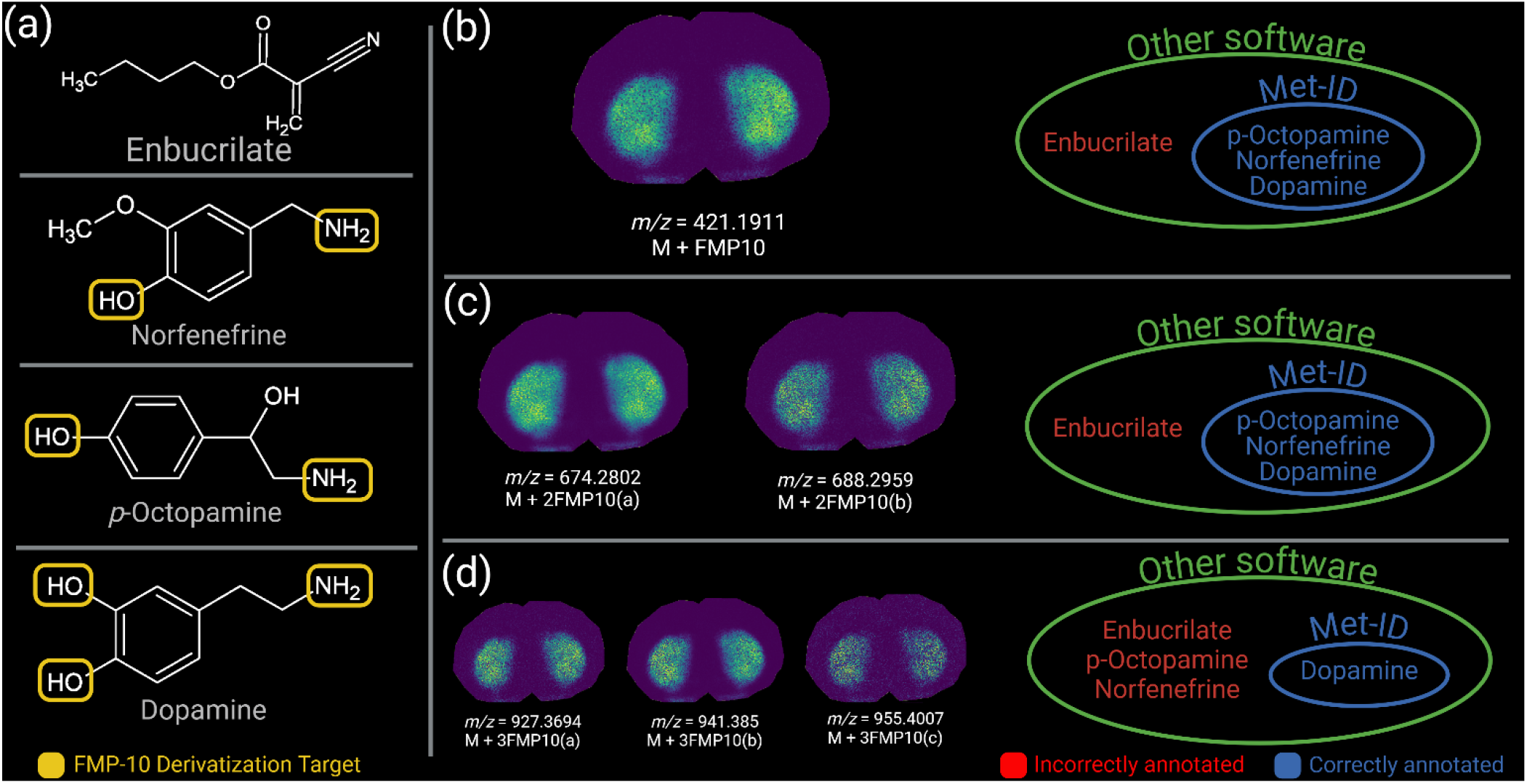
Comparing Met-ID to other annotation software. The four metabolites seen in (a) share the same molecular formula (C_8_H_11_NO_2_) and monoisotopic *m/z* ratio (153.07898). (b-d) shows how Met-ID annotations differ from other software for (b) single, (c)double, and (d) triple derivatized species. Met-ID, in contrast with other annotation software, takes information about derivatizing matrices into account and filters annotations based on molecular structures. FMP-10, which was the matrix used for this example, derivatizes primary amines and phenolic hydroxyls, highlighted in yellow in (a). As an example, only dopamine contains three derivatization targets, which is why it is the only annotation at, e.g., *m/z* 927.3694. Other software that do not consider chemical structures, would annotate all four molecules in (a). Furthermore, the consistent lateral distribution in the brain tissue section of the annotated ions (b-d) supports the conclusion that the identified species is dopamine. Ion images are shown using a rainbow scale (representing ion intensity scale) for clear visualization. Lateral resolution 100 μm.

Figure S1 shows how Met-ID can be used to discover new isomers of target metabolites

Dopamine can exist in two distinct derivatized forms when double derivatized with FMP-10. This occurs because one methyl group may be lost during the process, leading to a demethylated form (denoted as A), or it may remain intact, resulting in a dehydrogenated form (denoted as B) (Figure S2). Consequently, the derivatized dopamine produces characteristic peaks at *m/z* 421.1911 for dopamine 1A, *m/z* 674.2802 for dopamine 2A (demethylated) and *m/z* 688.2959 for dopamine 2B (dehydrogenated).

Moreover, in MSI experiments, a lock mass is often employed to ensure the mass spectrometer generates accurate *m/z* values. The choice of lock mass depends on several factors, including the chemical matrix used. The difference between the theoretical and experimental *m/z* values is zero at the lock mass, but deviations often increase for *m/z* ratios further away from the lock mass. To counteract this, Met-ID allows users to input known peaks across the spectrum to measure mass deviation and fit a correction curve. An example of this process is shown in Figure S3, where several known *m/z* values are provided. The calibration curve is fitted using nlopt’s,^18^ Newuoa^19^ and Bobyqa^20^ algorithms consecutively. The adjusted masses are then calculated using Equation 1 and can be considered more accurate, assuming the known peaks are correctly annotated.

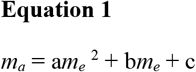

where *m*_*a*_ is the adjusted *m/z, m*_*e*_ is the experimental *m/z*, and a, b, c are fitted variables.

An experimental example was conducted using a dataset of 36 chemical standards, including catecholamines and amino acids, of which 28 were classified as endogenous in the HMDB. The dataset was analyzed using Fourier-transform ion cyclotron resonance (FTICR)-MSI. The SCiLS (Bruker Daltonics) T-ReX2 feature-finding algorithm, with 90% coverage and weak noise filtering settings, identified 256 *m/z* features, which were exported as a CSV file. Peaks with *m/z* values below that of FMP-10 (*m/z* 268.112076), along with peaks corresponding to FMP-10 and its fragments, were manually excluded, refining the dataset to 170 *m/z* features. This refined list was imported into Met-ID for further analysis (Figure 4a).

**Figure 4.**
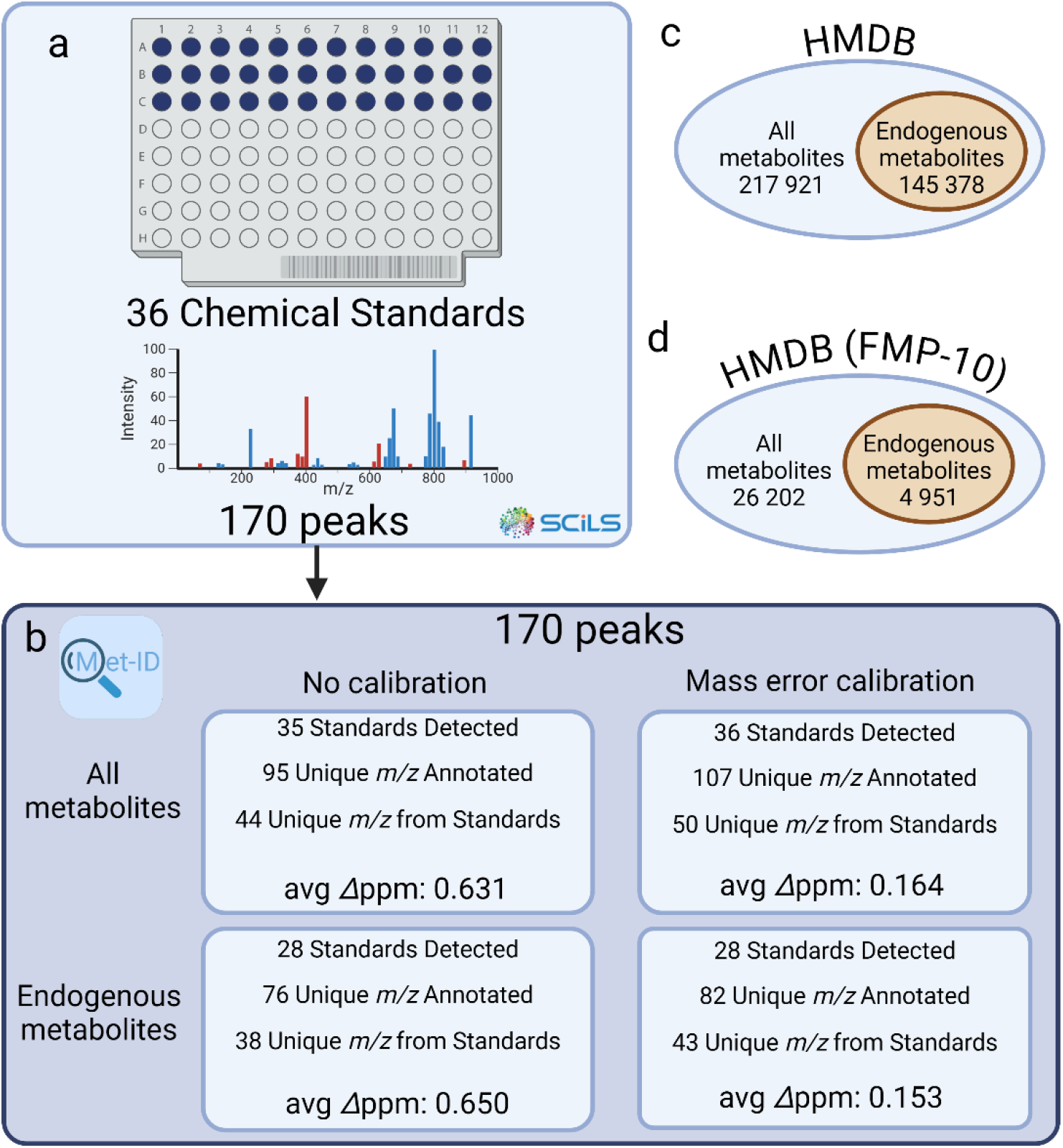
Met-ID analysis of chemical standards on a MALDI target plate. (a) A dataset of 36 chemical standards, including catecholamines and amino acids, generated a feature list of 256 *m*/*z* ratios using the SCiLS T-ReX2 algorithm. *m*/*z* ratios below that of FMP-10 and those corresponding to FMP-10 or its fragments were excluded, reducing the list to 170 *m*/*z* ratios. (b) Four searches were conducted using Met-ID: all metabolites in the HMDB with and without mass error calibration, and only endogenous metabolites with and without calibration. (c) The number of metabolites identified in the HMDB and the number of endogenous metabolites are shown. (d) Only metabolites derivatized by FMP-10 are included. Unidentified *m*/*z* ratios correspond to fragments outlined in Table S1.

Four distinct searches were performed in Met-ID using a 2 ppm identification window (Figure 4b). The first search targeted all metabolites potentially derivatized by FMP-10 (Figures 4c-d), resulting in 95 annotated *m/z* features (55.9%). Of these, 44 annotations corresponded to derivatives of 35 chemical standards. One standard fell outside the 2 ppm window and was not annotated, while multiple derivatives were observed for several standards. The remaining annotations corresponded to fragments of the chemical standards (Table S1).

Restricting the search to endogenous metabolites significantly reduced the number of potential candidates from 26,202 to 4,951 (Figure 4d). This search yielded 76 annotated *m/z* features (44.7% of the dataset). Of these, 38 (50%) were derivatives of the chemical standards, with 28 of the 36 standards annotated at least once. As expected, the remaining eight standards were not annotated as endogenous in the HMDB.

Next, mass error calibration was applied to the *m/z* values using a previously described method. This resulted in annotation of 107 *m/z* features (62.9%) for all metabolites and 82 *m/z* features (48.2%) for endogenous metabolites. The calibration was based on four dopamine peaks, two GABA peaks and a lock mass derived from FMP-10 (Table S2). Notably, all 36 chemical standards were annotated at least once when searching the entire HMDB, and all 28 endogenous standards were successfully identified in the restricted search.

The calibration process increased the number of annotated *m/z* features by 12.6% for all metabolites and by 7.9% for endogenous metabolites. This improvement was also reflected in the mean delta ppm between the observed and theoretical *m/z* values (excluding peaks used for calibration). Before calibration, the mean delta ppm for all metabolites was 0.631 ppm, which decreased by 74% to 0.164 ppm after calibration. The mean delta ppm for endogenous metabolites dropped from 0.650 ppm to 0.153 ppm, representing a 76.5% reduction.

As previously noted, the initial dataset of 256 *m/z* features was reduced to 170 after removing 43 peaks (16.8%) with *m/z* values below FMP-10 and 43 peaks (16.8%) identified as originating from FMP-10 or its fragments. These exclusions ensured that only relevant features were retained for further analysis.

This workflow demonstrates Met-ID’s capability to efficiently annotate metabolites in complex datasets using derivatization-specific filters and mass error calibration. By refining the input dataset and conducting targeted searches, Met-ID accurately identified metabolites derivatized by FMP-10, including multiple derivatives and endogenous compounds. The application of mass error calibration further enhanced the annotation accuracy, significantly reducing delta ppm values and increasing the number of reliable annotations. These results highlight Met-ID’s effectiveness in processing large datasets while minimizing false positives, making it a robust tool for both exploratory and routine metabolomics analyses.

In a subsequent example, we used data from a rat brain tissue section analyzed with a timsTOF fleX instrument (Bruker Daltonics) and FMP-10 as the reactive matrix. The T-ReX^2^ peak-picking algorithm in SCiLS was applied with 90% coverage and no noise filtering, detecting 576 *m/z* features. These features were exported as a CSV file and imported into Met-ID. Like in the example with chemical standards, Met-ID was run with a 2 ppm identification window. Searching for all metabolites potentially derivatized by FMP-10 resulted in annotation of 92 *m/z* features. After restricting the search to endogenous metabolites, 61 *m/z* features were annotated. The feature list was recalibrated using the same known peaks as in the chemical standards example, with the calibration details provided in Table S3. After recalibration, annotations were obtained for 90 and 59 *m/z* features for all metabolites and endogenous metabolites, respectively. Although the calibration resulted in two fewer annotations in both groups, the delta ppm decreased by 16.21% and 18.39% for all and endogenous metabolites, respectively (Table S4). This suggests that the reductions in annotations were likely due to the removal of false positives rather than true positives.

For comparison, another rat brain tissue section was analyzed using the conventional NEDC (N-(1-napthyl)ethylendiamine dihydrochloride) matrix. This analysis yielded 2,527 *m/z* features identified with the T-ReX^2^ feature-finding algorithm, applied with no spatial noise filtering and 95% coverage in SCiLS. These features were imported into Met-ID and searched against a database of endogenous metabolites, focusing on [M-H]^-^ and [M+Cl]^-^ adducts within a 2 ppm identification window. Initially, 445 *m/z* features (17.6%) were annotated. However, after applying Met-ID’s recalibration feature using 10 known peaks distributed across the spectrum (Table S5), the number of annotated features increased to 512 (20.3%), representing a 50% improvement in annotations. The mean delta ppm decreased by 13% when using Met-ID’s recalibration function.

A feature list containing 1538 *m/z* values was imported into Met-ID to benchmark performance. Using FMP-10 with a 2 ppm identification window, the software completed the identification process in approximately 1300 ms per attempt, measured from the moment the identify button was clicked to display the results. The benchmarking tests were carried out on a Windows desktop computer equipped with an Intel® Core™ i7-10700K CPU running at 3.80 Ghz. The performance was tested on a single thread.

### Annotating Chemically Derivatized Metabolites Using Met-ID and MS2

Several MS2 spectra were collected from FMP-10 derivatized chemical standards to construct a database for use in Met-ID. Currently, the MS2 database exclusively contains FMP-10 derivatized compounds. Since FMP-10 can produce multiple derivatized species depending on the functional groups present in the target molecules, double-derivatized fragments were also included in the MS2 data collection. The Met-ID MS2 database currently contains 154 derivatized forms originating from 66 chemical standards, including spectra at collision-induced dissociation (CID) ranges between 0-50 eV. Additionally, the MS2 database includes peaks from FMP-10 itself.

Met-ID functions as a frontend for viewing and searching these MS2 spectra, enabling targeted exploration. Users can search the database by molecular monoisotopic mass to locate specific product ions within the spectra. For instance, product ions specific to phenolic hydroxyl derivatization (*m/z* = 286.122641) and primary amine derivatization (*m/z* = 285.138625) are catalogued. Molecules that contain only phenolic hydroxyls, e.g., DOPAC, are identified by the phenolic hydroxyl-specific peak, whereas molecules that have only primary amines, e.g., spermidine, are identified by the primary amine-specific peaks. Molecules that contain both functional groups, e.g., dopamine, yield both product ion peaks. Figure S4 shows the product ion peaks from FMP-10 derivatization.

An additional feature of Met-ID is its ability to perform MS2 matching using cosine similarity^21-23^, a metric that quantifies the similarity between two spectra by treating them as vectors. A cosine similarity value of 1 indicates perfect spectral alignment, whereas a value of 0 indicates no similarity. In Met-ID, users can import MS2 spectra in mzML format and specify a bin size, which determines the resolution of the comparison. Cosine similarity is then calculated based on alignment of the intensities at matching *m/z* values within the specified bin size.

In Table 1, comparisons are made between several experimental MALDI-MS2 spectra and the Met-ID database. Experimental spectra obtained from tissue samples were compared with database spectra collected from derivatized standards on a MALDI-target plate. The corresponding MS2 spectra are presented in Figure S4.

**Table 1.** Comparison of MS2 from tissue to MS2 of standards in the Met-ID database. The names and adducts of the tissue metabolites are shown in the first two columns, followed by the collision induced dissociation (CID) and isolation window. The bin size is selected within Met-ID to decide the threshold defining what is considered the same. The best DB hit is the database MS2 spectrum with the highest cosine similarity to the MS2 spectrum queried. The rightmost column shows the cosine similarity between the queried spectra and the best matching database spectra. However, in the case of α-tocopherol, the correct hit (0.809) had a lower cosine similarity than that of the best match (0.865) (20 eV), which is why both numbers are shown.

| Tissue metabolite | Adduct | CID | Isolation window (Da) | Bin size | Best DB hit | Rank of correct hit | Cosine similarity of top hit & correct hit |
| --- | --- | --- | --- | --- | --- | --- | --- |
| Serotonin | 1A | 30 eV | 1 Da | 0.01 | Serotonin 1A 30 eV | 1st | 0.964 |
| DOPAC | 2A | 30 eV | 1 Da | 0.01 | DOPAC 2A 30 eV | 1st | 0.587 |
| Dopamine | 1A | 30 eV | 1 Da | 0.01 | Dopamine 1A 30 eV | 1st | 0.989 |
| Histidine | 1A | 30 eV | 1 Da | 0.01 | Histidine 1A 30 eV | 1st | 0.603 |
| Norepinephrine | 2A | 30 eV | 1 Da | 0.01 | Norepinephrine 2A 30 eV | 1st | 0.8 |
| Taurine | 1A | 20 eV | 1 Da | 0.01 | Taurine 1A 20 eV | 1st | 0.126 |
| α-Tocopherol | 1A | 30 eV | 1 Da | 0.01 | α-Tocopherol 1A 20 eV | 3rd | 0.865&0.809 |

In some cases, experimental spectra may extend beyond the precursor ion’s *m/z* ratio, introducing noise into the comparison. To address this, Met-ID allows users to define a maximum *m/z* value to be considered, typically slightly above the precursor ion’s *m/z* ratio. As demonstrated in Table S6, reanalysis of the spectra with a maximum *m/z* ratio threshold improves the specificity of the cosine similarity calculation. If no upper limit is defined, the calculation includes the entire spectrum.

The most common bin size used in the comparisons in Table 1 was 0.01 Da. Larger bin sizes, such as 1 Da, may reduce the specificity by grouping all intensities within a given range (e.g., *m/z* 421-422) as a single peak with a bin size of 1 Da. Conversely, smaller bin sizes require near-perfect mass accuracy between experimental and database spectra to achieve a high cosine similarity, making such comparisons more challenging. Table S7 demonstrates the effect of varying the bin size on the MS2 spectra of dopamine 1A and norepinephrine 2B.

### Met-ID Malleability

To enhance its versatility, Met-ID allows users to customize their own versions. FMP-10 is a suitable matrix for experiments targeting neurotransmitters, and the base version of Met-ID is optimized for its use. However, Met-ID also provides tools for calculating and supporting the different masses added by other derivatizing matrices. Additionally, since some metabolites are not included in the HMDB, Met-ID enables users to add new entries, ensuring the software remains adaptable to specific research needs.

In MS2 mode, users have two options for data input: comparing data to the existing database or adding new spectra to the database. The MS2 database is built using spectra imported in the mzML format, a widely accepted open standard for mass spectrometry data. The base version of Met-ID comes preloaded with several MS2 spectra derived from chemical standards. It also allows users to expand the database by adding their own MS2 spectra. Furthermore, Met-ID provides tools for editing the database, enabling users to remove entries if errors occur during data input, ensuring database integrity.

In summary, evaluation of Met-ID across diverse datasets demonstrates its capability to address key challenges in metabolite identification for MALDI-MSI. Its workflows enable efficient data import, accurate annotation of derivatized metabolites and enhanced precision through recalibration. Specifically, Met-ID has the following attributes: (a) Met-ID uniquely incorporates derivatization-specific filtering, enabling the accurate identification of metabolites with functional groups targeted by agents like FMP-10. (b) The combination of mass matching and spectral comparison, i.e., dual MS1 and MS2 functionality, ensures comprehensive coverage of metabolite annotations. (c) Recalibration significantly reduces mass errors and improves confidence in the results, as evidenced by higher annotation rates and lower delta ppm values. (d) Met-ID efficiently processes large datasets, making it suitable for high-throughput spatial metabolomics. These features underscore Met-ID’s role as a robust and versatile tool for spatial metabolomics, offering both customization and broad applicability for diverse research needs.

## DISCUSSION

Met-ID represents a significant advancement in metabolite identification for MALDI-MSI, particularly when using derivatizing matrices like FMP-10. Although Met-ID has been demonstrated to perform well with MSI data, it is equally applicable to non-imaging datasets, broadening its usability. Traditional metabolomics software often struggles with the accurate annotation of derivatized metabolites, as they typically do not account for chemical structures, derivatization processes or the possibility of multiple derivatizations per target molecule. Met-ID addresses these challenges through a dynamic, customizable database and enhanced MS1 and MS2 functionalities. By utilizing the chemical properties and structural information of analytes, Met-ID reduces false positives and improves the confidence and accuracy of metabolite identifications. Additionally, incorporation of advanced MS2 matching algorithms, including cosine similarity, enables more precise comparison between user data and established databases.

One of Met-ID’s key strengths lies in its flexibility, allowing users to customize the software by adding new derivatizing matrices, functional groups or metabolites not present in existing databases such as the HMDB. This adaptability extends its applicability beyond neurotransmitter analysis, for which it was initially developed, enabling its use in a wide range of metabolomics studies.

Another software commonly used in spatial metabolomics is Metaspace, which has become a popular tool for automatic metabolite annotation in MSI datasets^12^. Although Metaspace is a widely used platform for spatial metabolomics offering robust annotation for high-resolution MSI data, it was originally developed for underivatized datasets and later adapted for chemically derivatized MSI data^24^. Metaspace provides automatic annotation and supports custom databases, but it has limitations in addressing the complexities of derivatized datasets, particularly when multiple derivatized products or specific MS2 features are involved.

In contrast, Met-ID has been specifically designed for derivatized MSI data and incorporates detailed knowledge of derivatization reactions, such as functional group specificity (e.g., phenolic hydroxyls and primary amines), and the ability to account for multiple derivatization states. These features allow Met-ID to accurately annotate complex patterns of derivatization, which Metaspace cannot do without significant manual intervention.

Our results demonstrate the effectiveness of Met-ID across various datasets, including chemical standards and biological samples such as brain tissue sections. The software’s ability to recalibrate based on known peaks and its support for custom MS2 spectra makes it a powerful tool for routine and exploratory mass spectrometry analyses. For instance, Met-ID can distinguish between dopamine derivatives that retain or lose a methyl group based on their MS2 fragmentation patterns, a level of precision not directly supported by Metaspace. These unique features make Met-ID an essential tool for applications such as neurotransmitter mapping and other studies requiring detailed structural insights. However, some limitations exist, particularly regarding the software’s pre-configured finite number of MS2 spectra. The introduction of automated MS2 collection for MSI would be a natural progression to address this limitation.

Met-ID’s design is in alignment with the FAIR (Findable, Accessible, Interoperable, and Reusable) principles, ensuring broad usability and supporting future development. As novel identification methods and tools emerge, Met-ID will evolve, especially with the open-source community’s contributions. The software’s modularity and adaptability, combined with its advanced identification capabilities, suggest that Met-ID has the potential to become a valuable tool in spatial metabolomics research.

In summary, we present Met-ID, a software solution for metabolite identification specifically tailored to MALDI-MSI, capable of generating matching metabolite lists within seconds. As the FMP-10 derivatizing matrix was developed in-house, its datasets were used as benchmarks during the software’s development. Met-ID utilizes derivatization chemistry and MS2 spectral comparisons to effectively annotate metabolites that are difficult for conventional tools to identify. The ability to recalibrate mass errors enhances confidence in metabolite identification, reduces false positives and supports robust downstream analyses.

The complete codebase is freely available on GitHub, allowing users to install and customize their own versions. Contributions through pull requests are encouraged to keep the software up-to-date with emerging identification methods. Met-ID also offers extensive customization for non-programmers, enabling users to easily add or remove metabolites, functional groups and matrices directly within the software. The tool’s modular design and open-source community support provide a strong foundation for future enhancements.

## METHODS

### Programming Languages

Met-ID was built in the Tauri software framework, which utilizes a Rust programming backend with a general web frontend consisting of HTML, CSS and Typescript. The code was written in Visual Studio Code with the rust-analyzer extension as a tool for writing Rust. The code was compiled into executable files for Windows, MacOS and Debian systems through Github Actions, allowing for simple compilations upon changes to the codebase.

Met-ID is also dependent on rdkit^25^, a C++ framework for computational chemistry. The installation of rdkit is slightly more difficult than an average PYPI project but often works with anaconda. The rdkit functionality was written in python files and turned into executable files using PyInstaller. The executable files were then added to the Tauri project using what Tauri calls “Sidecar”.

### Met-ID Modularity

Met-ID was designed to be modular and was thus built by separating the code for MS1 from that of MS2. This was to avoid system wide bugs, as well as making it easier to make changes to the codebase.

### Met-ID Data Analysis

Met-ID is based on the openly available Human Metabolome DataBase (HMDB) and Lipidmaps and contains files parsing the HMDB^14^ and Lipidmaps^16^ into SQLite databases that Met-ID can read. These databases can easily be directly interacted with within Met-ID or outside of Met-ID using any method of interacting with SQLite databases. The user can manually update the database. Pull requests are highly encouraged to help improve Met-ID, as well as adding support.

### MS2 Database

A library of chemical standards were analyzed in MS2. The MS2 analyses used a 1 Da isolation window and spectra collected at different collision energies (0-50 eV). Different collision energies were used to discriminate the peak in question from neighboring peaks.

### FAIR Software

Met-ID was developed according to the FAIR Guiding Principles for software.

## ASSOCIATED CONTENT

### Supporting Information

The Supporting Information is available free of charge at https://pubs.acs.org/doi/XXXXXXXXXXXX.

Isomers of dopamine, derivatives of FMP-10, product ions specific to FMP-10 derivatization of different functional groups., mirrored MS2 spectra comparing tissue metabolites to Met-ID database spectra, list of FMP-10 chemical standards and number of fragments. list of FMP-10 *m*/*z* ratios used in calibration of an MS1 experiment on chemical standards, list of FMP-10 *m*/*z* ratios used in calibration of an MS1 experiment on tissue, comparison of delta ppm error between FMP-10 searches on tissue data, list of negative mode *m*/*z* ratios used in calibration of an MS1 experiment, comparison of Met-ID database MS2 and tissue MS2 with maximum *m/z*, comparison of Met-ID database MS2 and tissue MS2 with different bin sizes, HTX sprayer methods, MALDI-MSI of murine tissue with FMP-10, MALDI-MSI of rat tissue in negative mode, MALDI-MSI of chemical standards with FMP-10 (PDF).

## Code Availability

All code as well as executable files for Windows, MacOS and Debian systems is publicly available on Github via *https://github.com/pbjarterot/Met-ID*

## Author Information

**Authors**

**Patrik Bjärterot** - Department of Pharmaceutical Biosciences, Spatial Mass Spectrometry, Science for Life Laboratory, Uppsala University, Uppsala SE-75124, Sweden

**Anna Nilsson** - Department of Pharmaceutical Biosciences, Spatial Mass Spectrometry, Science for Life Laboratory, Uppsala University, Uppsala SE-75124, Sweden

**Reza Shariatgorji** - Department of Pharmaceutical Biosciences, Spatial Mass Spectrometry, Science for Life Laboratory, Uppsala University, Uppsala SE-75124, Sweden

**Theodosia Vallianatou** - Department of Pharmaceutical Biosciences, Spatial Mass Spectrometry, Science for Life Laboratory, Uppsala University, Uppsala SE-75124, Sweden

**Ibrahim Kaya** - Department of Pharmaceutical Biosciences, Spatial Mass Spectrometry, Science for Life Laboratory, Uppsala University, Uppsala SE-75124, Sweden

**Per Svenningsson** - Department of Clinical Neuroscience, Karolinska Institutet, Stockholm, SE-17177, Sweden

**Lukas Käll** - Science for Life Laboratory, Department of Gene Technology, KTH — Royal Institute of Technology, Solna SE-17165, Sweden

## Author Contributions

P.B: conceptualization, investigation, formal analysis, methodology, validation, writing original draft, and review and editing. A.N, R.S, T.V, I.K: methodology, validation, and review and editing. P.S: conceptualization, resources, writing, review and editing, and funding acquisition. LK: methodology, validation, supervision, review and editing. P.E.A.: conceptualization, methodology, resources, validation, writing, review and editing, visualization, project administration, supervision, and funding acquisition.

## Notes

A.N., R. S. and P.E.A. are co-founders and shareholders in Tag-ON AB and are holders of the patent ‘Reactive desorption and/or laser ablation ionization matrices and use thereof’, no. PCT/SE2019/050197. The other authors declare that they have no known competing interests.

## ACKNOWLEDGMENTS

This work was supported by the Swedish Research Council (grants 2022-04198 and 2021-03293), the Swedish Brain Foundation (grant FO2023-0241), and the Science for Life Laboratory.

