## Supplemental Information for "Met-ID: An Open-Source Software for Comprehensive Annotation of Multiple On-Tissue Chemical Modifications in MALDI-MSI"

| SUPPORTING FIGURES | | Page S2-S6 |
|---|---|---|
| Figure S1 | Isomers of dopamine. | Page S2 |
| Figure S2 | Derivatives of FMP-10. | Page S2 |
| Figure S3 | Product ions specific to FMP-10 derivatization of different functional groups. | Page S3 |
| Figure S4 | Mirrored MS2 spectra comparing tissue metabolites to Met-ID database spectra. | Page S4-S6 |

| SUPPORTING TABLES | | Page S7-S10 |
|---|---|---|
| Table S1 | List of FMP-10 chemical standards and number of fragments. | Page S7 |
| Table S2 | List of FMP-10 *m/z* ratios used in calibration of an MS1 experiment on chemical standards. | Page S7 |
| Table S3 | List of FMP-10 *m/z* ratios used in calibration of an MS1 experiment on tissue. | Page S8 |
| Table S4 | Comparison of delta ppm error between FMP-10 searches on tissue data. | Page S8 |
| Table S5 | List of negative mode *m/z* ratios used in calibration of an MS1 experiment. | Page S9 |
| Table S6 | Comparison of Met-ID database MS2 and tissue MS2 with maximum *m/z*. | Page S9 |
| Table S7 | Comparison of Met-ID database MS2 and tissue MS2 with different bin sizes. | Page S10 |

| SUPPORTING METHODS | | Page S11 |
|---|---|---|
| Method S1 | HTX sprayer methods. | Page S11 |
| Method S2 | MALDI-MSI of murine tissue with FMP-10. | Page S11 |
| Method S3 | MALDI-MSI of rat tissue in negative mode. | Page S11 |
| Method S4 | MALDI-MSI of chemical standards with FMP-10 | Page S11 |

### Supporting Figures

421.1911 | Show only identified | Search...

| Observed m/z | Adjusted m/z | Matched name | Adduct | Formula | Matched delta mass (Da) | Matched difference (ppm) | Matched theoretical mass | MSMS available? | Coverage |
|---|---|---|---|---|---|---|---|---|---|
| 421.191100 | 421.191100 | 2-(2-Amino-1-hydroxyethyl)phenol | [M+FMP10] | C8H11NO2 | 4.54020e-5 | 0.107794 | 421.191055 | Not available | 1 / 3 |
| 421.191100 | 421.191100 | Norfenefrine | [M+FMP10] | C8H11NO2 | 4.54020e-5 | 0.107794 | 421.191055 | Not available | 1 / 3 |
| 421.191100 | 421.191100 | Dopamine | [M+FMP10] | C8H11NO2 | 4.53990e-5 | 0.107787 | 421.191055 | Available | 1 / 6 |
| 421.191100 | 421.191100 | p-Octopamine | [M+FMP10] | C8H11NO2 | 4.53990e-5 | 0.107787 | 421.191055 | Available | 1 / 3 |
| 421.191100 | 421.191100 | Vanillylamine | [M+FMP10] | C8H11NO2 | 4.53990e-5 | 0.107787 | 421.191055 | Not available | 1 / 3 |

**Figure S1. Isomers of dopamine.** Met-ID shows all isomers of dopamine that may be derivatized by FMP-10. In this example, the *m/z* value of single derivatized dopamine was queried. Met-ID displayed all the isomers, as well as which metabolites have MS2 spectra collected in the database.

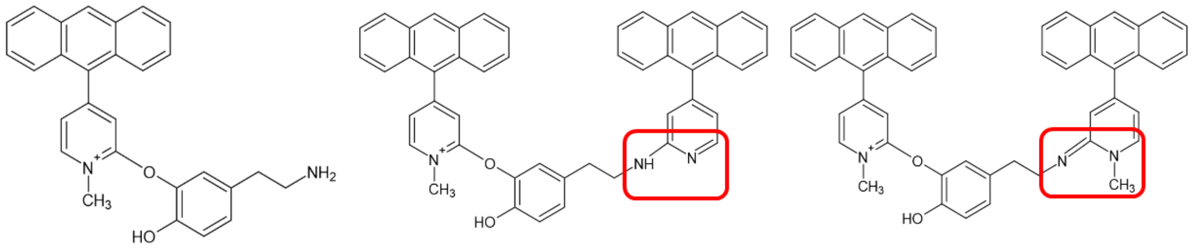

(a) Dopamine 1A *m/z* 421.1911

(b) Dopamine 2A *m/z* 674.2802

(c) Dopamine 2B *m/z* 688.2959

**Figure S2. Derivatives of FMP-10.** (a) Single-derivatized dopamine (dopamine 1A), and (b) double-derivatized dopamine (dopamine 2A). As highlighted by the red box, the FMP-10 molecule in dopamine 2A loses a methyl group, resulting in the loss of one positive charge and giving the molecule an overall charge of +1. (c) Double-derivatized dopamine (Dopamine 2B) retains the methyl group but loses a hydrogen. This causes a double bond to shift from the aromatic ring to the dopamine nitrogen. Consequently, the FMP-10 nitrogen loses its positive charge, resulting in an overall charge of +1. Both A and B derivatives can form if the target molecule undergoes double derivatization. Triple derivatization follows a similar pattern, with the addition of a C derivative (e.g., dopamine 3A, dopamine 3B and dopamine 3C). The letters assigned to these derivatives correspond to their *m/z* ratios, with A representing the lowest *m/z* ratio and B and C representing progressively higher *m/z* ratios for double- and triple-derivatized molecules, respectively.

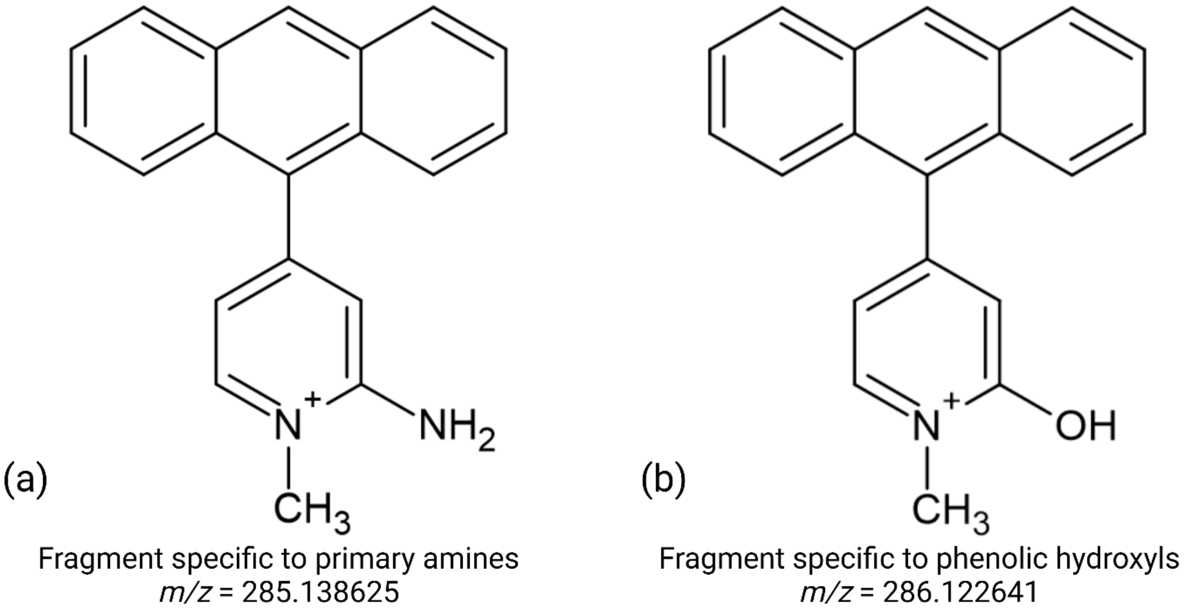

**Figure S3. Product ions specific to FMP-10 derivatization of different functional groups.** (a) Fragment specific to primary amine derivatization. (b) Fragment specific to phenolic hydroxyl derivatization.

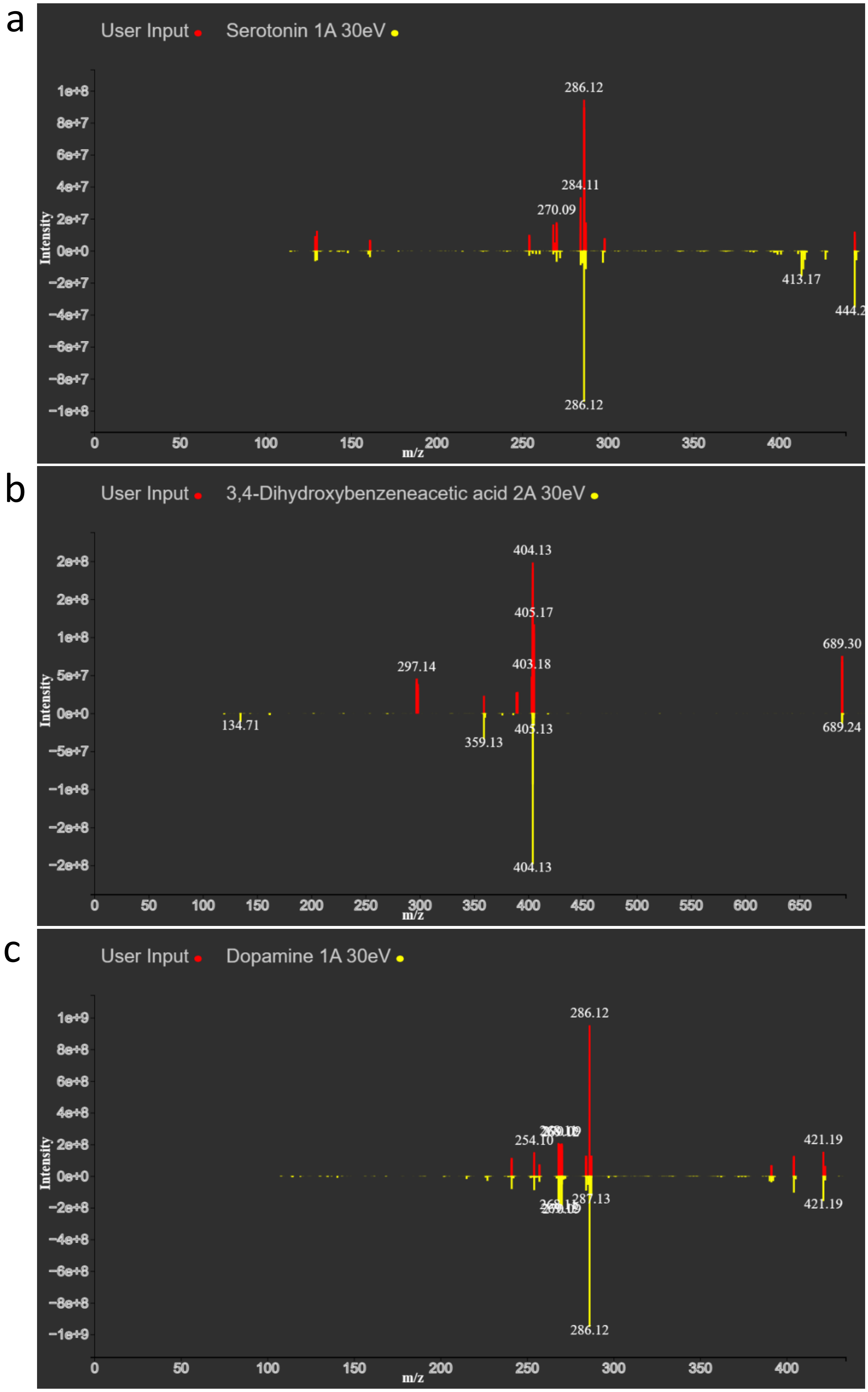
a
User Input
Serotonin 1A 30eV
Intensity
m/z
286.12
284.11
270.09
413.17
444.2
286.12
b
User Input
3,4-Dihydroxybenzeneacetic acid 2A 30eV
Intensity
m/z
404.13
405.17
403.18
297.14
689.30
134.71
359.13
405.13
689.24
404.13
c
User Input
Dopamine 1A 30eV
Intensity
m/z
286.12
254.10
421.19
287.13
421.19
286.12

d

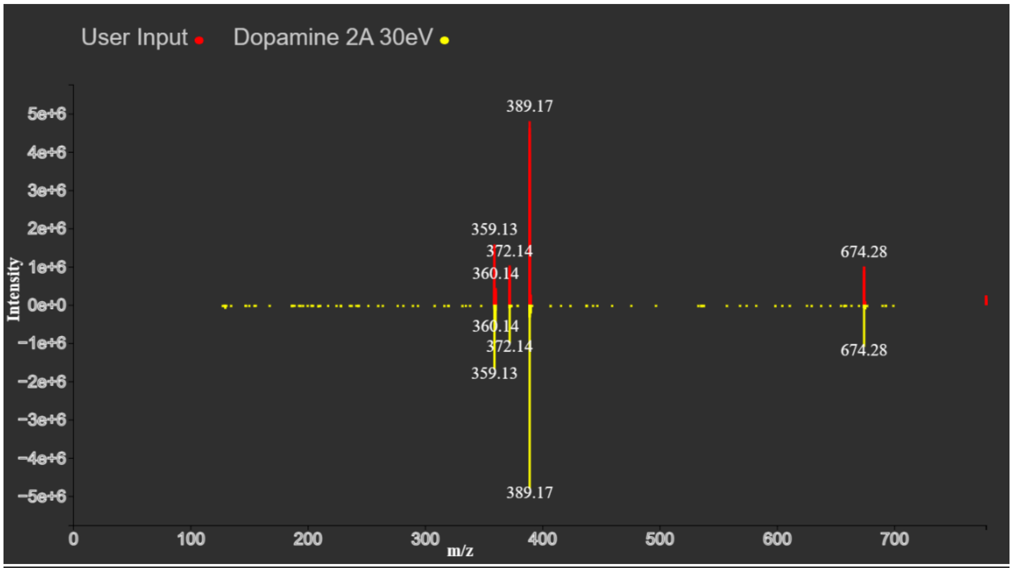
User Input
Dopamine 2A 30eV
389.17
359.13
372.14
360.14
674.28
360.14
372.14
359.13
674.28
389.17
5e+6
4e+6
3e+6
2e+6
1e+6
0e+0
−1e+6
−2e+6
−3e+6
−4e+6
−5e+6
Intensity
0
100
200
300
400
500
600
700
m/z

e

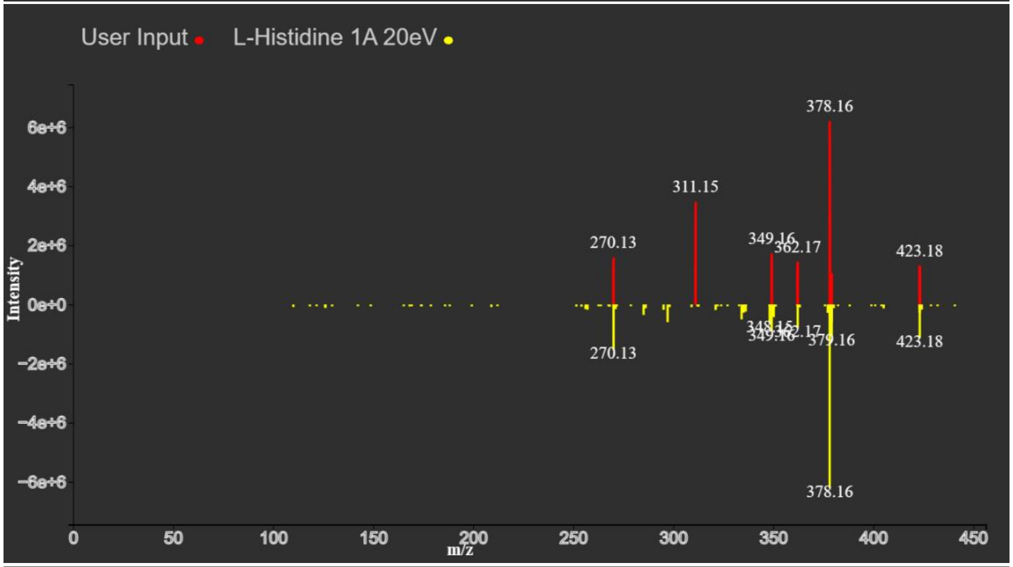
User Input
L-Histidine 1A 20eV
378.16
311.15
270.13
349.16
362.17
423.18
270.13
379.16
423.18
378.16
6e+6
4e+6
2e+6
0e+0
−2e+6
−4e+6
−6e+6
Intensity
0
50
100
150
200
250
300
350
400
450
m/z

f

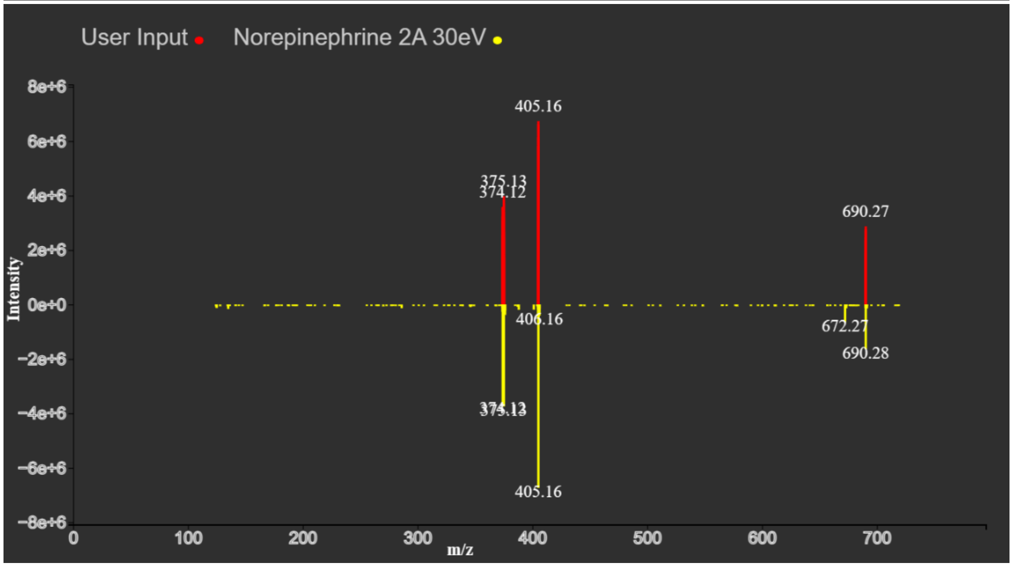
User Input
Norepinephrine 2A 30eV
405.16
375.13
374.12
690.27
406.16
672.27
690.28
374.12
375.13
405.16
8e+6
6e+6
4e+6
2e+6
0e+0
−2e+6
−4e+6
−6e+6
−8e+6
Intensity
0
100
200
300
400
500
600
700
m/z

g

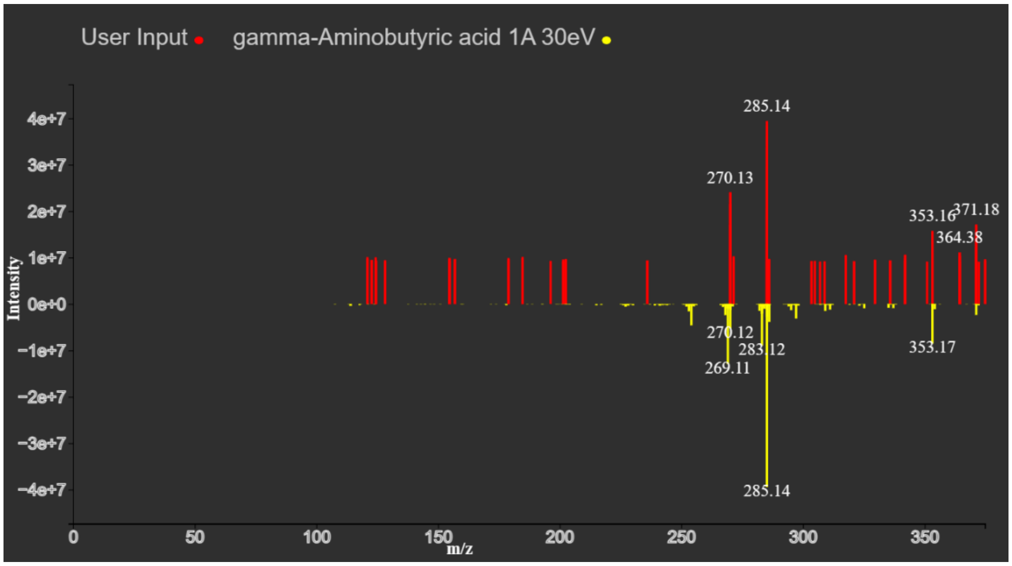

**Figure S4. Mirrored MS2 spectra comparing tissue metabolites to Met-ID database spectra.** The mirrored spectra are as follows (a) serotonin 1A 30 eV, (b) DOPAC 2A 30 eV, (c) dopamine 1A 30 eV, (d) dopamine 2A 30 eV, (e) histidine 1A 20 eV, (f) norepinephrine 1A 30 eV, and (g) GABA 1A 30 eV. The labeling of derivatives is explained in the caption of Figure S2.

### Supporting Tables

**Table S1. List of FMP-10 chemical standards and their fragments.** Certain chemical standards tend to fragment more frequently, sometimes producing fragments with *m*/*z* ratios similar to those of other metabolites or fragments. These fragments can generate significant peaks that may be included in the *m*/*z* feature lists, potentially leading to false positives. For example, homovanillic acid contributed to 10 out of the 170 peaks in the *m*/*z* feature list but was correctly identified only once (as homovanillic acid 1A).

| Chemical standard | Number of fragments | Chemical standard | Number of fragments |
|---|---|---|---|
| 5-Hydroxyindoleacetic acid | 17 | Norepinephrine | 4 |
| 3,4-Dihydroxyphenylglycol | 14 | Vanillylmandelic acid | 4 |
| Homovanillic acid | 10 | Metyrosine | 3 |
| Vanillactic acid | 10 | Histamine | 3 |
| L-Lysine | 9 | Adenine | 3 |
| L-Histidine | 9 | L-Tyrosine | 3 |
| L-Arginine | 9 | L-Dopa | 3 |
| Epinephrine | 9 | Normetanephrine | 2 |
| Serotonin | 9 | *p*-Octopamine | 2 |
| 3,4-Dihydroxybenzeneacetic acid (DOPAC) | 8 | Putrescine | 2 |
| Norsalsolinol | 7 | Spermidine | 2 |
| 3-Methoxytyrosine | 7 | 5,6-Dihydroxyindole | 2 |
| 3-Methoxytyramine | 7 | Melevodopa | 1 |
| Carbidopa | 7 | γ-Aminobutyric acid | 1 |
| Salsolinol | 6 | Spermine | 1 |
| Dopamine | 5 | Tryptamine | 1 |
| Hordenine | 5 | 4-Hydroxy-benzaldehyde | 1 |
| Tyramine | 4 | L-Phenylalanine | 0 |

**Table S2. List of FMP-10 *m/z* ratios used in calibration of an MS1 experiment on chemical standards.** A curve was fitted to this data and used to recalibrate the input data to more closely match the theoretical *m*/z ratio.

| Metabolite | Derivative (*) | Experimental *m/z* | Theoretical *m/z* | ppm difference |
|---|---|---|---|---|
| Dopamine | 1A | 421.19144 | 421.191055 | 0.91 |
| Dopamine | 2A | 674.27992 | 674.280203 | -0.42 |
| Dopamine | 2B | 688.29543 | 688.295854 | -0.62 |
| Dopamine | 3B | 941.38171 | 941.385004 | -3.5 |
| Lock mass | | 555.22310 | 555.223104 | 0 |
| GABA-H2O | 1A | 353.16516 | 353.164840 | 0.91 |
| GABA | 1A | 371.17562 | 371.175405 | 0.58 |

(*) Derivative labels are as described in the caption of Figure S2.

**Table S3. List of FMP-10 *m/z* ratios used in calibration of an MS1 experiment on tissue.** A curve was fitted to this data and used to recalibrate the input data to more closely match the theoretical *m/z*.

| Metabolite | Derivative (*) | Experimental *m/z* | Theoretical *m/z* | ppm difference |
|---|---|---|---|---|
| Dopamine | 1A | 421.191 | 421.191055 | -0.13 |
| Dopamine | 2A | 674.2802 | 674.280203 | 0 |
| Dopamine | 2B | 688.2954 | 688.295854 | -0.66 |
| Dopamine | 3B | 941.3851 | 941.385004 | 0.10 |
| Lock mass | | 555.2231 | 555.2231 | 0 |
| GABA-H2O | 1A | 353.1648 | 353.16484 | -0.11 |
| GABA | 1A | 371.1753 | 371.175405 | -0.28 |

(*) Derivative labels are as described in the caption of Figure S2.

**Table S4. Comparison of delta ppm error between FMP-10 searches on tissue data.** The ppm difference between the theoretical and observed *m/z* for all *m/z* features with annotations for the four different searches. The *m/z* features used in the calibration were excluded.

| | Endogenous metabolites (Δppm) | All metabolites (Δppm) |
|---|---|---|
| No mass error calibration | 0.524 | 0.519 |
| Mass error calibration | 0.427 | 0.435 |
| Reduction (%) | 18.39 | 16.21 |

**Table S5. List of negative mode *m/z* ratios used in calibration of an MS1 experiment.** A curve was fitted to this data and used to recalibrate the input data to more closely match the theoretical *m/z*.

| Metabolite | Adduct | Experimental *m/z* | Theoretical *m/z* | ppm difference |
|---|---|---|---|---|
| Aspartic acid | $[M-H]^-$ | 132.03033 | 132.030231 | 0.75 |
| Glucose | $[M+Cl]^-$ | 215.03293 | 215.032789 | 0.66 |
| Glutamine | $[M-H]^-$ | 145.06182 | 145.061866 | -0.32 |
| Glutathione | $[M-H]^-$ | 306.07669 | 306.07653 | 0.52 |
| Taurine | $[M-H]^-$ | 124.00746 | 124.007388 | 0.58 |
| LysoPE(22:6) | $[M-H]^-$ | 524.27837 | 524.278263 | 0.20 |
| PE(38:4) | $[M-H]^-$ | 766.53898 | 766.539229 | -0.32 |
| PE(40:6) | $[M-H]^-$ | 790.53881 | 790.539229 | -0.53 |
| PI(38:4) | $[M-H]^-$ | 885.55043 | 885.549859 | 0.64 |
| Ascorbic acid | $[M-H]^-$ | 175.02491 | 175.024812 | 0.56 |

**Table S6. Comparison of MS2 from tissue to MS2 of standards in the Met-ID database with the maximum *m/z* ratio limit.** This table is similar to Table 1 with set maximum *m/z* ratio considered. Some MS2 spectra have collected data greater than the precursor ion mass, which impacts the cosine similarity. Notably, DOPAC showed a considerable increase in cosine similarity when disallowing higher *m/z* values from being considered.

| Tissue metabolite | Derivative (*) | CID | Isolation window (Da) | Max *m/z* | Bin size (Da) | Best DB hit | Rank of correct hit | Cosine similarity of top hit & correct hit |
|---|---|---|---|---|---|---|---|---|
| Dopamine | 1A | 30 eV | 1 Da | 422 | 1 | Dopamine 1A 30 eV | 1st | 0.989 |
| Dopamine | 1A | 30 eV | 1 Da | 422 | 0.1 | Dopamine 1A 30 eV | 1st | 0.989 |
| Dopamine | 1A | 30 eV | 1 Da | 422 | 0.01 | Dopamine 1A 30 eV | 1st | 0.99 |
| Dopamine | 1A | 30 eV | 1 Da | 422 | 0.001 | Dopamine 1A 30 eV | 1st | 0.794 |
| Dopamine | 1A | 30 eV | 1 Da | 422 | 0.0001 | DHPME 1A 30 eV | >20th | 0.405&0.049 |
| Norepinephrine | 2A | 30 eV | 1 Da | 691 | 1 | Norepinephrine 2A 30 eV | 1st | 0.983 |
| Norepinephrine | 2A | 30 eV | 1 Da | 691 | 0.1 | Norepinephrine 2A 30 eV | 1st | 0.982 |
| Norepinephrine | 2A | 30 eV | 1 Da | 691 | 0.01 | Norepinephrine 2A 30 eV | 1st | 0.87 |
| Norepinephrine | 2A | 30 eV | 1 Da | 691 | 0.001 | Norepinephrine 2A 30 eV | 1st | 0.642 |
| Norepinephrine | 2A | 30 eV | 1 Da | 691 | 0.0001 | Epinephrine 2B 40 eV | 5th | 0.168&0.072 |

(*) Derivative labels are as described in the caption of Figure S2.

**Table S7. Comparison of MS2 from tissue to MS2 of standards in the Met-ID database with different bin sizes.** Dopamine and norepinephrine are compared to show the effect of changing bin sizes. In both cases, a bin size of 0.01 Da gave the highest cosine similarity. While larger bin sizes gave similar cosine similarities, the specificity decreased. The correct hit for serotonin with a 1 Da bin size had the 10th highest cosine similarity. However, decreasing the bin size reduced the sensitivity, and thus the cosine similarity, due to small calibration changes, which may occur if spectra are collected at different times.

| **Tissue metabolite** | **Derivative (*)** | **CID** | **Isolation window (Da)** | **Max *m/z*** | **Bin size (Da)** | **Best DB hit** | **Rank of correct hit** | **Cosine similarity of top hit & correct hit** |
|---|---|---|---|---|---|---|---|---|
| Serotonin | 1A | 30 eV | 1 Da | 445 | 0.01 | Serotonin 1A 40 eV | 10th | 0.952&917 |
| DOPAC | 2A | 30 eV | 1 Da | 690 | 0.01 | DOPAC 2A 30 eV | 1st | 0.798 |
| Dopamine | 1A | 30 eV | 1 Da | 422 | 0.01 | Dopamine 1A 30 eV | 1st | 0.99 |
| Dopamine | 2A | 30 eV | 1 Da | 675 | 0.01 | Dopamine 2A 30 eV | 1st | 0.987 |
| Histidine | 1A | 30 eV | 1 Da | 424 | 0.01 | Histidine 1A 30 eV | 1st | 0.738 |
| Norepinephrine | 2A | 30 eV | 1 Da | 691 | 0.01 | Norepinephrine 2A 30 eV | 1st | 0.87 |
| GABA | 1A | 30 eV | 2 Da | 372 | 0.01 | Putrescine 1A 20 eV | 2nd | 0.75&0.724 |

(*) Derivative labels are as described in the caption of Figure S2.

### Supporting Methods

#### Method S1. HTX sprayer methods

*N*-(1-naphthyl) ethylenediamine dihydrochloride (NEDC) was prepared at a concentration of 7 mg/ml in 70 % MeOH and sonicated in an ultrasonic bath for 30 s. The matrix solvent was sprayed over the sample and the dried standard spots using a TM-Sprayer (HTX Technologies) in 14 passes with a 0.07 ml/min flow of pushing solvent (50% ACN), nozzle temperature of 50°C, track spacing of 2 mm, nozzle velocity of 1100 mm/min and $N_2$ gas pressure of 6 psi. The derivatizing matrix FMP-10 was used according to a previously described protocol[5]. In brief, 20 passes of 4.4 mM FMP-10 in 70% acetonitrile were sprayed (nitrogen pressure of 6 psi) in horizontal lines over the tissue with a solvent flow rate of 80 µL/min, nozzle temperature of 90 °C, nozzle velocity of 1100 mm/min and track spacing of 2 mm.

#### Method S2. MALDI-MSI of murine tissue with FMP-10

Mass spectrometry imaging data of tissue sections coated with FMP-10 were acquired at 50 µm lateral resolution on a timsTOF flex MALDI-2 mass spectrometer (Bruker Daltonics). A single focus beam was used to collect 450 shots/pixel at 10 kHz. The scan range was set to 300 – 1000 *m/z* with TIMS on the $1/K_0$ range of 0.7-1.65 and a ramp time of 300 ms. The method was externally calibrated using red phosphorous and internally calibrated using the matrix peak at *m/z* 569.2224. The T-ReX$^2$ peak-picking algorithm in SCiLS (Bruker Daltonics) was used with 90% coverage and no noise filtering.

#### Method S3. MALDI-MSI of rat tissue in negative mode

A MALDI-FTICR-MS instrument (Solarix XR 7T-2Ω, Bruker Daltonics, Bremen) was used for acquiring data from the NEDC-coated sections. The fourier transform ion cyclotron resonance (FTICR) scan range was set to 86-1000 *m/z* at 100 µm lateral resolution, summing up 100 shots/pixel. The laser focus was set to a frequency of 1 kHz. The quadrupole isolation (Q1) *m/z* was set to 120 and the transfer optics and time-of-flight (TOF) was set to 4 MHz and 0.650, respectively. No lock mass was used. The methods were externally calibrated using red phosphorous. The T-ReX$^2$ feature-finding algorithm with no spatial noise filtering and 95% coverage was used to find features in SCiLS (Bruker Daltonics).

#### Method S4. MALDI-MSI of chemical standards with FMP-10

The same MALDI-FTICR-MS instrument was used for acquiring data from the chemical standards coated with FMP-10. The scan range was set to 150.50-1050 *m/z* at 200 µm lateral resolution and 100 shots per pixel. The laser focus was set to a frequency of 1 kHz. The quadrupole isolation (Q1) *m/z* was set to 379 and the transfer optics and TOF were set to 4 MHz and 0.750 ms, respectively. A FMP-10 derived peak at *m/z* 555.2231 was used as a lock mass for internal *m/z* calibration. Feature finding was performed in SCiLS (Bruker Daltonics) using the T-ReX$^2$ feature finding algorithm with 90% coverage and weak noise filtering settings.
